# Tropomyosin 1 promotes platelet adhesion and clot contraction separate from its roles in developmental hematopoiesis

**DOI:** 10.1101/2025.07.31.667883

**Authors:** Po-Lun Kung, Victor Tsao, Alina D Peshkova, Oscar A. Marcos-Contreras, Kim Ha, Gennadiy Fonar, Nkemdilim Okoli, Brian M Dulmovits, Rong Qiu, Rolf D Bates, Janelle Yeboah, Carson Shalaby, Tyler Truex, Soomin Jeong, Vladimir R Muzykantov, Jacob W Myerson, Christopher S Thom

## Abstract

Genome-wide associations studies (GWAS) have linked the *Tropomyosin 1* (*Tpm1*) gene locus to quantitative blood trait variation, but related mechanisms are unclear. *Tpm1* encodes an actin-binding protein that stabilizes actin filaments and influences cell adhesion, signaling, and actomyosin contractility. Murine *Tpm1* deficiency enhances embryonic hemogenic endothelial cell specification, but it was unclear if these effects extended to postnatal hematopoiesis. We used *Cdh5*^*Cre*^ or *Vav*^*Cre*^ models to conditionally ablate *Tpm1* in endothelium or hematopoietic cells. Both models produced knockout mice in normal Mendelian ratios with complete *Tpm1* ablation in postnatal blood. Endothelial *Tpm1* deletion increased hemogenic endothelial cell specification, but did not change hematopoietic progenitor cell production nor adult blood counts. This suggested separate roles for *Tpm1* in the embryonic and adult blood systems. GWAS suggested genetic architecture specifically linking decreased *TPM1* expression to increased platelet count. We examined platelet lifespan and function to explain these findings. *Tpm1KO* increased platelet lifespan and diminished adhesion to fibronectin and fibrinogen. Decreased platelet clearance could explain increased platelet count in GWAS. Platelet fibrin binding is necessary for blood clot contraction, which reduces vascular occlusion following initial hemostasis. *Tpm1KO* reduced clot contraction and enhanced clot formation with worsened vascular occlusion in a ferric chloride-induced stroke model. These findings reveal a new role for *Tpm1* in platelet function, offering insight into how cytoskeletal regulation impacts human platelet traits and pointing to novel targets to modify stroke risk and thrombotic disease.

## Introduction

Genome-wide associations studies (GWAS) have linked thousands of loci to quantitative blood trait variation, but related genes and mechanisms remain elusive for most sites^1,2^. Platelet traits, such as platelet count (PLT) and mean platelet volume (MPV), are highly heritable (∼54-87%)^3^. Yet mechanisms and cell types that are relevant for altered platelet traits are complex. Factors intrinsic to developing blood cells (hematopoiesis), megakaryocytes (megakaryopoiesis), or platelets (thrombopoiesis) can influence platelet formation, Systemic factors like obesity or inflammation - extrinsic to developing blood cells - also influence platelet traits via influences on the bone marrow microenvironment or mature platelet lifespan in circulation^4^. Blood cells interact with all organs and impact complex disease states, highlighting a need to reveal mechanisms underlying blood trait variation^5^.

We identified the *Tropomyosin 1 (TPM1)* gene locus from blood trait GWAS^6^. Tpm1 encodes an actin-binding protein that regulates actin cytoskeletal dynamics in many cell types^7^. Actin broadly impacts cell structure, movement, and signaling. *TPM1* deficiency increases the formation of hemogenic endothelial cells (HEC), the embryonic precursors to hematopoietic stem and progenitor cells (HSPCs) that ultimately colonize the bone marrow and support lifelong hematopoiesis^8^ (**Fig. 1**). Embryonic lethality from *Tpm1*-related cardiac dysmorphology prevented assessment of how *Tpm1* knockout impacted postnatal hematopoiesis and blood cell function^8,9^.

**Figure 1.**
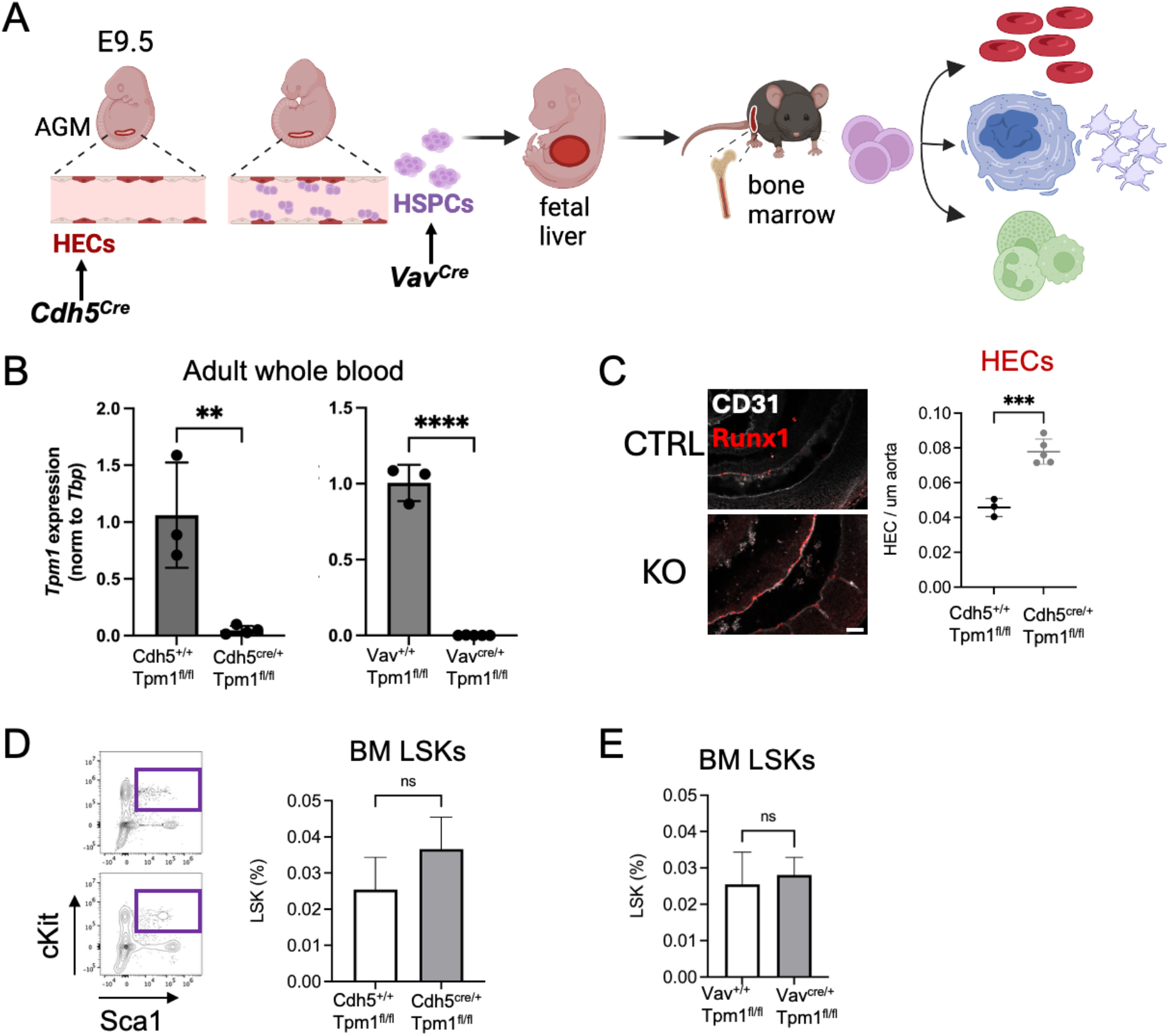
Tropomyosin 1 deficiency impacts formation of hemogenic endothelium but not postnatal bone marrow HSPC quantity,. A. Schematic of *Cdh5*^*Cre*^ and *Vav*^*Cre*^ models that are induced at specific stages of developmental hematopoiesis. B. *Tpm1* mRNA is eliminated in adult peripheral blood cells following conditional embryonic deletion. Semiquantitative real time PCR results normalized to *TBP*. C. Embryonic *Tpm1* deletion in endothelial cells increases the incidence of Runx1 HECs. D. Embryonic *Tpm1* deletion with *Cdh5*^*Cre*^ does not increase bone marrow LSK abundance. E. *Tpm1* deletion in embryonic HSPCs with *Vav*^*Cre*^ does not change bone marrow LSK abundance. ^****^p<0.0001, ^***^p<0.001, ^**^p<0.01, ns, not significant.

The strongest GWAS signals at the *TPM1* gene locus relate to platelet traits (p<10^−100^)^1,2^. Platelets produced from megakaryocytes are critical for hemostasis and inflammation^10^. Blood vessel injury exposes factors that cause platelet activation and adherence to the site, initiating hemostatic responses^11^. Aggregated platelets form a temporary clot and entrap erythrocytes and white blood cells to enhance plug formation and inflammatory responses. These processes prevent hemorrhage and initiate wound healing, but large clots can obstruct blood flow and risk embolization^11^. Clot contraction is necessary to stabilize clots and permit resumption of blood flow. Clot formation demands platelet activation, fibrin binding, and actin-mediated cytoskeletal remodeling^12,13^. Therefore, effective hemostasis requires platelet activation, adhesion, cytoskeletal remodeling – all of which are actin-based properties to varying degrees.

We hypothesized that *Tpm1* impacted platelet function via actin regulation. We designed this study ascertain if the effects of *Tpm1* perturbation during embryonic life carried through to postnatal hematopoiesis, or if *Tpm1* has separable effects on platelet function. Using conditional knockout mice, we reveal novel roles for *Tpm1* in platelet adhesion, clot formation and thrombosis that are distinct from effects on embryonic hematopoiesis. Our findings advance our understanding of how cytoskeletal regulatory factors impact hemostasis.

## Results

### Tpm1 has separable roles in embryonic and postnatal hematopoiesis

To circumvent cardiac-related embryonic lethality from *Tpm1* deficiency^8,9^, we generated conditional *Tpm1* knockout (*KO*) models with a floxed *Tpm1* exon 3 (*Tpm1*^*fl*^)^14^. Exon 3 is present in all known *Tpm1* isoforms. We crossed *Tpm1*^*fl*^ mice with *Cdh5*^*Cre*^ or *Vav*^*Cre*^ mice to abrogate *Tpm1* expression in endothelial and nascent hematopoietic stem and progenitor cells (HSPCs), respectively^15,16^. We used *Cdh5*^*Cre*^-mediated deletion evaluate the effects of *Tpm1* deletion that occurred at different stages of developmental hematopoiesis (**Fig. 1A**). The *Cdh5*^*Cre*^ construct is active in HE cells^15^, which we presumed would result in deletion in all HSPCs and peripheral blood cells through adulthood. The *Vav*^*Cre*^ allele can efficiently abrogate gene expression in HSPCs and peripheral blood through postnatal life^16,17^. Both models are well validated in the context of developmental hematopoiesis. We validated efficient recombination and abrogation of *Tpm1* mRNA in peripheral mononuclear blood cells in adult mice compared to littermate controls in both *Cdh5*^*Cre*^ and *Vav*^*Cre*^ models (**Fig. 1B**).

We further validated by whole mount immunostaining that E9.5 *Cdh5*^*Cre*^ *Tpm1*^*fl/fl*^ embryos exhibited increased Runx1^+^ HE cell production at E9.5 compared with littermate controls (**Fig. 1C**), similar to *Tpm1-*deficient mouse embryos^8^. *Cdh5*^*Cre*^ *Tpm1*^*fl/fl*^ and *Vav*^*Cre*^ *Tpm1*^*fl/fl*^ pups were born at expected Mendelian ratios and exhibited normal viability into adulthood.

We next asked if increased HE cell production in *Cdh5*^*Cre*^ *Tpm1*^*fl/fl*^ embryos conferred an increase in HSPCs in postnatal bone marrow hematopoiesis (**Fig. 1A**). By flow cytometry, we observed no differences in Lin^-^cKit^+^Sca1^+^ bone marrow HSPCs or peripheral blood counts at steady state (**Fig. 1D** and **Supplementary Table S1**). We similarly found no significant differences in adult bone marrow of *Vav*^*Cre*^ *Tpm1*^*fl/fl*^ mice compared to littermate controls, despite previously confirmed Cre activation during fetal life^16^ (**Fig. 1E** and **Supplementary Table S2**). The lack of change in the postnatal blood progenitors after Tpm1 deletion led us to conclude that the effects of *Tpm1* deficiency during embryonic hematopoiesis separate from any effects on postnatal hematopoiesis.

### Human genetics implicates Tpm1 in mature platelet function

Human genome wide association study (GWAS) data have linked single nucleotide polymorphisms at the *TPM1* gene locus with quantitative blood trait variation^1,2^. In light of the disconnect between fetal hematopoietic biology and postnatal bone marrow traits, these GWAS signals suggested that *TPM1* had some role in hematopoietic stem and progenitor cells and/or mature blood cells in circulation. The most statistically significant associations are with platelet traits, including platelet count (PLT) and mean platelet volume (MPV, **Fig. 2A**). However, there are also genome-wide significant signal associations with hemoglobin (HGB, **Fig. 2A**). There is also sub-genome-wide significant signal for variation in red blood cell (RBC) and white blood cell count (WBC, **Fig. 2A**).

**Figure 2.**
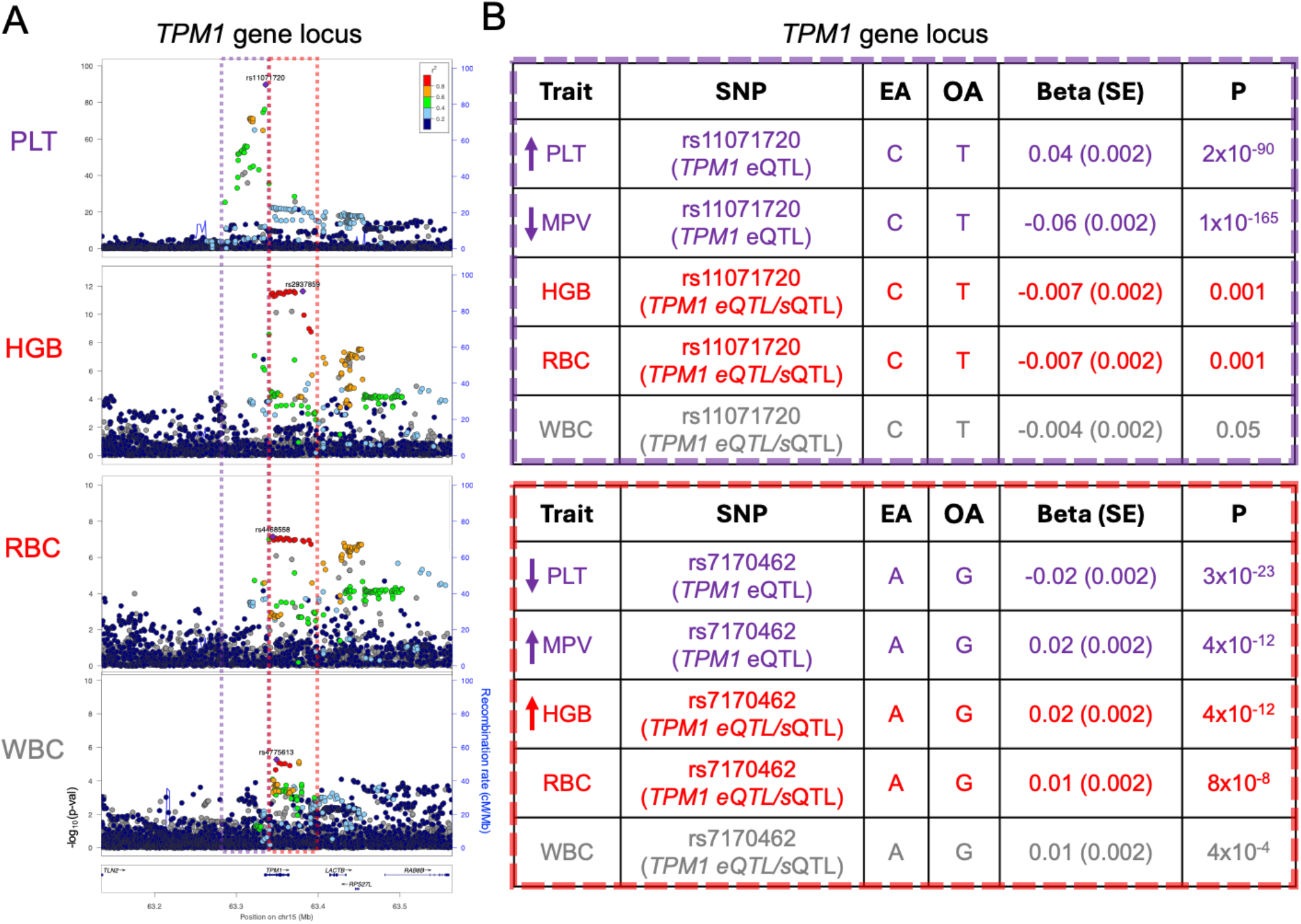
Genetic colocalization analysis points to separate genetic signals for altered platelet count vs erythroid and white blood cell parameters. A. Locus zoom plots suggest separate genetic signals for platelet traits and erythroid traits at the *TPM1* gene locus. Purple box indicates loci with statistically significant signals for increased platelet count. Red box indicates loci with significant signals for erythroid traits. B. Key single nucleotide polymorphisms (SNPs) at the *TPM1* gene locus that impact blood traits in human GWAS. SNPs that act as quantitative trait loci for *TPM1* expression (eQTL) and/or splice variation (sQTL) are indicated. Effect alleles (EA) and other alleles (OA) for the indicated directional effects are shown.

Our prior work revealed that systemic factors like obesity can have coordinate impacts on blood traits across lineages, reflecting perturbations in HSPC metabolism and development^4,5^. In other cases, GWAS signals can have lineage-specific effects more likely to reflect impacts on mature blood cells in circulation. For example, genetically influenced lipid traits and tobacco use have erythroid-specific effects that suggest impacts on peripheral erythrocytes^18^. To discern if *TPM1* locus-related GWAS effects indicated coordinate effects across blood lineages, we performed genetic colocalization analysis^19^. Comparisons of PLT, HGB, and WBC effects revealed platelet-specific effects at the *TPM1* gene promoter and transcriptional start site that did not colocalize with erythroid or white cell traits (PP4 < 0.02, **Fig. 2B**). These sites include expression quantitative trait loci (eQTL) that overlie relevant hematopoietic transcription factor binding sites and alter *TPM1* expression^6^. Erythroid and white cell traits shared colocalized signal within the gene body (PP4 = 0.94, **Fig. 2B**). SNPs in this region include splice quantitative trait loci (sQTL) that alter splicing between *TPM1* exons without altering total *TPM1* mRNA levels^20^. These colocalization findings indicate separate lineage-specific effects on mature platelets and erythrocytes, or late-stage precursors for these blood cells, rather than effects on common progenitors for multiple lineages (e.g., HSPCs).

### Tpm1 impacts murine platelet lifespan and adhesion capabilities

We next chose to interrogate *TPM1* effects on platelet biology, given that these were statistically the strongest GWAS effects. SNPs that decrease *TPM1* expression are linked with increased PLT count (**Fig. 2A**). We reasoned that *TPM1* deficiency could either increase platelet formation and/or decrease platelet clearance in circulation. We have previously shown using in vitro models that *TPM1KO* megakaryocyte formation and maturation is normal^6^. Thus, we asked if platelet clearance was increased in *Vav*^*Cre*^ *Tpm1*^*fl/fl*^ mice, which could help explain an increased steady state platelet count. To elucidate the lifespan of platelets lacking *Tpm1*, we injected mice with anti-platelet CD42c^DyLt488^ antibody and monitored the lifespan of labelled CD42c^DyLt488+^/CD41^APC+^ platelets over time by flow cytometry. *Tpm1KO* mice had longer platelet lifespan compared to littermate controls, as evidenced by an increased area under the curve (5366±153 KO vs 4741±148 Ctrl, p<0.0001) and a ∼50% increased half-life (86 h vs 58 h in littermate controls by linear regression, **Fig. 3A**). Thus, *Tpm1* normally limits platelet lifespan in murine circulation.

**Figure 3.**
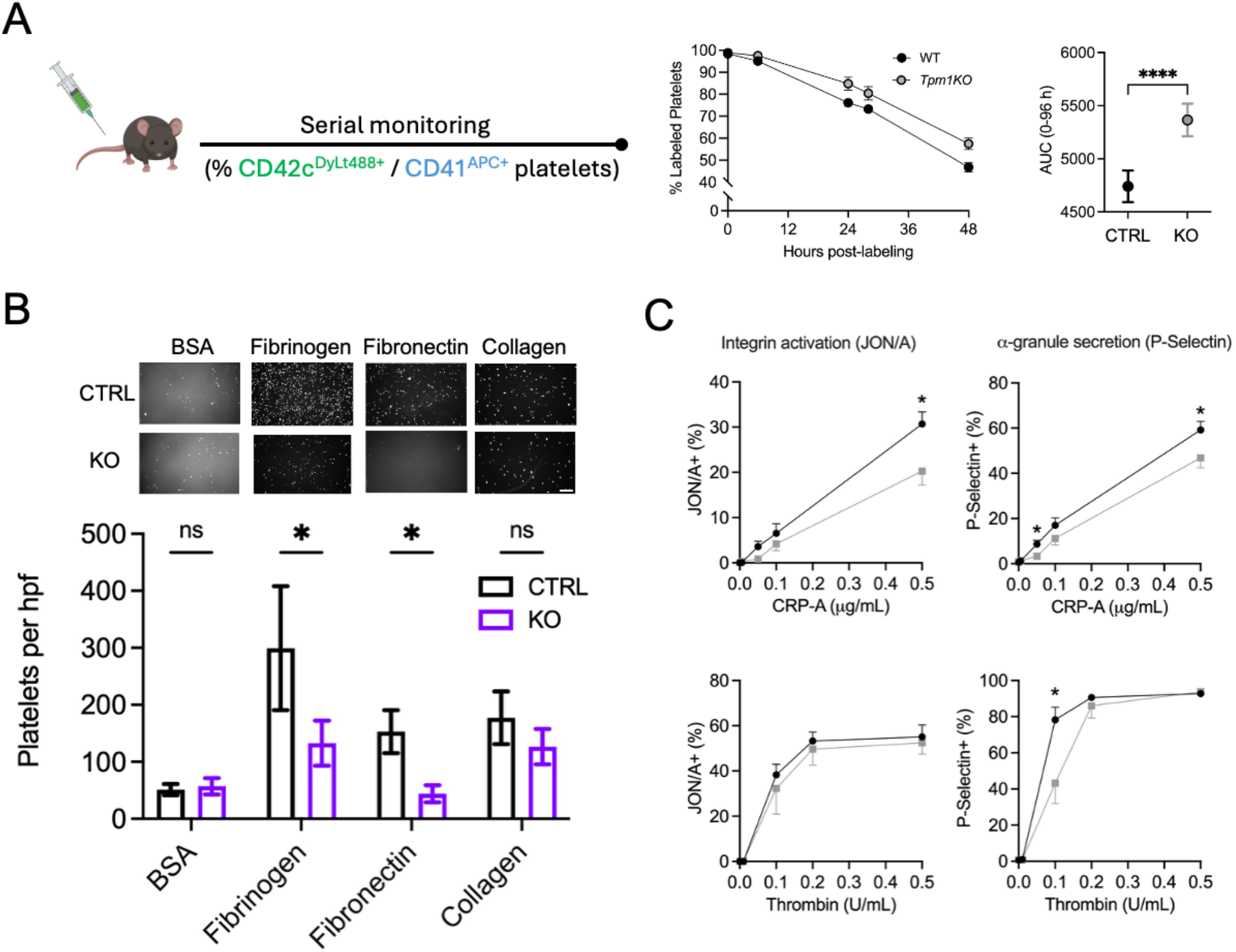
*Tpm1* knockout prolongs platelet lifespan and limits platelet adhesion. A. Platelet half-life measurement after labeling indicates longer time in circulation for *Tpm1KO* platelets. Platelets were labeled with anti-CD42c^DyLight488^ antibody and the percentage of platelets was measured by flow cytometry over time. B. Static focal adhesion experiments show decreased *Tpm1KO* platelet adhesion to Fibrinogen and Fibronectin-coated coverslips compared to wild type controls, but no significant change in adhesion to collage-coated coverslips (CTRL n=6-8, KO n=4-8 per substrate). Scale bar, 25 um. C. *Tpm1KO* platelets show variably reduced activation in response to agonists. ^*^p<0.05. ^****^p<0.0001.

We hypothesized that *Tpm1* deficiency could increase platelet lifespan by limiting clearance from circulation, via limiting adhesion to vascular walls and/or decreasing activation potential. Focal adhesion biology directly impacts platelet clearance by regulating interactions with vascular walls^21,22^. Focal adhesions contain nanoscale layers that are functionally specified by tropomyosin isoforms^23^. Tpm1 regulates adhesion maturation and disassembly in these structures^23^. To test *Tpm1KO* platelet adhesion, we compared adhesion of platelets from *Vav*^*Cre*^ *Tpm1*^*fl/fl*^ mice vs littermate controls in static cell adhesion assays using fibrinogen, fibronectin, or collagen substrates. These substrates directly bind and activate platelets through dedicated platelet membrane receptor protein complexes and can lead to adhesion and activation^22^. After incubating blood with matrix-coated coverslips for 30 min, we found more platelets retained on all substrate-coated cover slips compared with bovine serum albumin-coated controls (**Fig. 3B**). *Tpm1KO* compromised platelet adhesion to fibrinogen and fibronectin, and to a lesser (non-significant) effect on collagen (**Fig. 3B**).

We next evaluated murine platelet activation potential ex vivo using primary murine platelets from *Vav*^*Cre*^ *Tpm1*^*fl/fl*^ mice or littermate controls^24^. We measured integrin αIIbβ3 activation (CD41^+^JON/A-PE^+^) and degranulation (CD41^+^P-Selectin^+^) after incubation with thrombin, collagen-related peptide A (CRP-A), and adenosine diphosphate (ADP) agonists. *Tpm1KO* platelets showed mild changes in activation potential that were inconsistent across agonists (**Fig. 3C**). Our findings suggest that platelet *Tpm1KO* activation may be altered in certain contexts, but that *Tpm1* exerts more prominent effects on platelet adhesion properties.

### Tpm1 deficiency impairs clot contraction

We then sought to determine the functional consequences of altered platelet properties in *Tpm1KO* platelets. Hemostasis involves both clot formation and subsequent contraction, which together produce a stable hemostatic plug that stops bleeding effectively without obstructing blood flow^11,12,25^. Clot contraction is driven by activated platelets that adhere to fibrin fibers, forming a three-dimensional viscoelastic scaffold. One of the downstream effects of platelet activation is the interaction between intracellular actin and non-muscle Myosin IIA^26^.

We tested *Tpm1KO* platelet functionality using a blood clot contraction assay, which quantifies the degree of clot volume reduction as a measure of actomyosin-dependent platelet contractility^27–29^ (**Fig. 4A**). ^12,13,30,29,30^Compared to littermate controls, whole blood samples from *Tpm1KO* mice showed a significant delay in the initiation of clot contraction and decreased area under the curve for total clot contraction, while retaining normal contraction velocity and extent of maximal contraction (**Fig. 4B** and **Supplementary Fig. 1A**).

**Figure 4.**
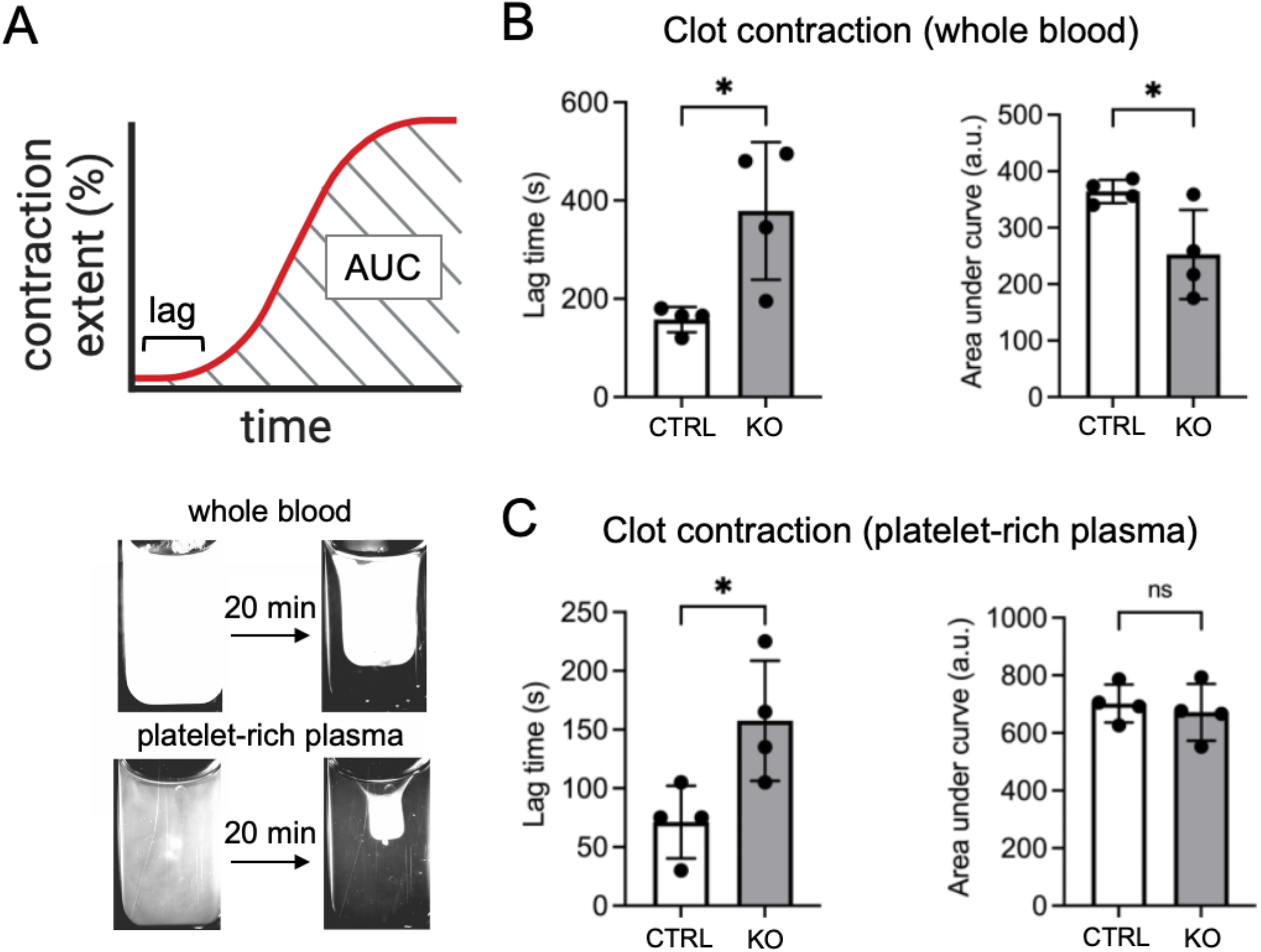
*Tpm1KO* delays initiation of clot contraction. A. Clot contraction assay schematic. B,. Clot contraction initiation is delayed and the area under the curve for clot contraction is limited in *Tpm1KO* whole blood compared to littermate controls. C. Clot contraction is delayed in *Tpm1KO* platelet-rich plasma (PRP) compared to littermate controls. ^*^p<0.05. ns, not significant.

We then determined if these findings were due to platelet-specific effects. Using platelet-rich plasma isolated from *Tpm1KO* and littermate controls, we observed a significant delay in the initiation of clot contraction lag time (**Fig. 4C** and **Supplementary Fig. 1B**). These results confirm a functional consequence for abnormal platelet function in the context of *Tpm1* deficiency, which is consistent with ex vivo functional readouts and associated with potential physiologic implications.

### Tpm1KO accentuates murine stroke formation and severity

Platelet focal adhesion is necessary for efficient clot contraction and prevention of pathologic thrombotic extension^30^. These effects are somewhat counterintuitive, since defective focal adhesion or platelet activation can also compromise hemostasis and result in bleeding^31,32^. To define the hemostatic implications for altered focal adhesion in *Tpm1* deficiency, we tested the effects of *Tpm1KO* in an established murine stroke model^33^. After ferric chloride-induced injury to the middle cerebral artery, we monitored blood flow and subsequent clot pathology. Compared to littermate controls, *Tpm1KO* shortened the time to abrogation of blood flow at the injury site (flow rate x time area under the curve [AUC] 30.1±5.1 vs 13.4±3.6, mean±SD, p=0.01, **Fig. 5A-B**).

**Figure 5.**
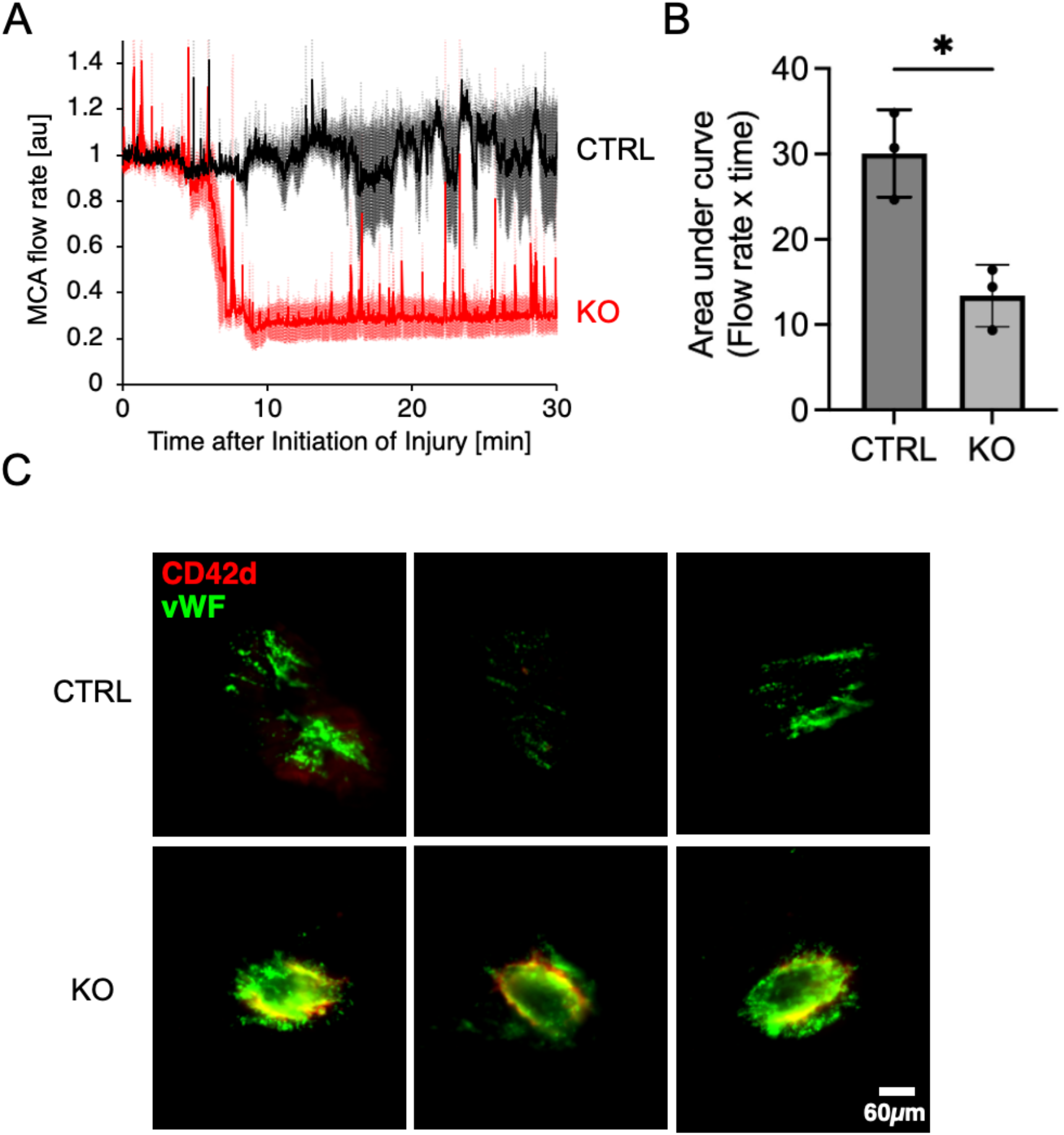
*Tpm1KO* enhances ferric chloride-induced middle cerebral artery (MCA) occlusion via enhanced platelet interactions with vWF. A. Flow rate curves depicting time to MCA occlusion (n=3 per genotype). B. Quantitative changes for MCA occlusion area under the curve (n=3 per genotype). ^*^p<0.05. C. Clot histology showing platelets (CD42d^+^) and vWF aggregation at the site of MCA vessel injury. KO mice showed more circumferential platelet aggregation and vWF accumulation.

MCA blood clot histology from control and KO mice confirmed an enhanced interaction of *TPM1KO* platelets with von Willebrand factor (vWF) in the context of vessel injury (**Fig. 5C**). However, we did not detect significant changes in bleeding times nor coagulation pathway activation at steady state in *TPM1KO* mice compared to littermate controls (**Supplementary Fig. 2**). These findings demonstrate that in the context of ferric chloride-induced vessel injury and inflammation, *TPM1KO* platelets have an enhanced response to promote blood clotting. These results provide a novel functional consequence for *Tpm1KO* on platelet biology and hemostasis.

## Discussion

This study reveals a novel role for *TPM1* in platelet adhesion and clot contraction. These effects appear distinct from *TPM1* impacts on embryonic hematopoiesis^8^ (**Fig. 1**) and are intrinsic to platelets, as opposed to endothelial cells or non-hematopoietic cell types that support vascular niches during embryonic development^34^. Our findings agree with emerging roles for *TPM1* in mediating focal adhesion stability^23^ and actomyosin contractility during clot contraction^11,12^. To our knowledge, *TPM1* is the first actin regulatory molecule implicated in clot contraction outside of Myosin IIA and fibrin^11,12^. We anticipate that our findings will lead to further insights into platelet biology and hemostasis, given the extensive prior work on *TPM1* in the context of cardiac sarcomeres^9^.

Mammalian actin filament diversity is achieved through expression of 4 genes (*TPM1-*4) that encode more than 40 isoforms through alternative splicing^7^. Each isoform can impart unique properties on actin filaments. Our murine and iPSC models disrupt all *TPM1* isoforms^6,8,35^ so we cannot yet parse isoform-specific *TPM1* functions in platelets. In fact, high molecular weight and low molecular weight *TPM1* isoforms have distinct actin regulatory roles in some cell types^36^. Future isoform-specific perturbation will parse individual tropomyosin contributions to actin cytoskeletal regulation of platelet functions. In addition, *TPM4* is highly expressed in platelets and *TPM4* loss causes macrothrombocytopenia (i.e., reducing platelet counts) with a mild effect on platelet function^37^. *TPM1* and *TPM4* may genetically interact to regulate distinct but complementary aspects of platelet biology, which could be elucidated through combined perturbation using genetic models.

Our findings can help explain GWAS signal at the *TPM1* gene locus^1,2^. Rather than effects on platelet production^37^, we propose that SNPs at the *TPM1* locus extend platelet lifespan to increase steady-state platelet count. We suspect that species-specific differences or environmental variations may underlie the normal platelet counts in *Tpm1KO* mice. Human exposure to infections, inflammation, and other stressors could amplify functional consequences of altered human platelet lifespan, enhancing the impact of allelic variation that changes *TPM1* expression (**Fig. 2**). The controlled living conditions of our inbred mice may mask phenotypic variation that emerges in real world human cohorts with large sample sizes. Future studies will explore the functional consequences of recurrent infection or inflammation on platelet biology in the context of *Tpm1* deficiency.

TPM1 likely facilitates platelet adhesion by stabilizing actin filaments at focal adhesion sites, possibly through binding and/or stabilizing Talin and integrin complexes at a nanoscale level^23^. Interestingly, Tpm1 cardiac pathology is ameliorated with concurrent Talin mutation^38^. This genetic interaction in cardiac dysfunction may extend to platelets. Our static adhesion results indicate that *Tpm1* loss perturbs platelet adhesion to fibrinogen and fibronectin, but not collagen (**Fig. 3**). Talin facilitates platelet adhesion to fibrinogen and fibronectin, while collagen binding can be Talin-independent through GPVI^22^. Thus, our findings agree with a molecular role for Tpm1 in conjunction with Talin at the platelet cell surface.

Our findings add a novel actin regulatory protein (TPM1) to the small group of molecules known to impair clot contraction and enhance thrombosis, which also includes fibrinogen, integrins, protease-activated receptors (PARs), and myosin IIA^11^. Defective clot contraction results in larger, less dense, mechanically fragile thrombi with increased risks of embolization and vascular occlusion^11,30^. Our findings highlight the role of actomyosin regulation in clot contraction, an established role for *Tpm1* in cardiac muscle ^*7*^. We expect *Tpm1* is relevant in platelets and potentially other cell types (e.g., erythrocytes^39^), since we identified defective clot contraction in assays of *Tpm1KO* whole blood but not PRP (**Fig. 4**). Promoting *Tpm1* activity or otherwise targeting related mechanisms offers a novel translational strategy to prevent or ameliorate thrombosis and stroke pathology.

This study supports the potential translational utility linked to defining genetic determinants of platelet and blood trait variation. Our findings define a novel factor (*Tpm1*) governing platelet adhesion, clot contraction, and thrombosis. Further work to establish developmental and postnatal roles of *Tpm1* will undoubtedly reveal links between actomyosin contractility, cytoskeletal regulation, and hemostasis.

## Materials and Methods

### Mouse model derivation and validation

All mouse experiments were approved by the Children’s Hospital Institutional Animal Care and Use Committee (IACUC). The floxed *Tpm1* allele was derived from the EUCOMM *Tpm1* GeneTrap-Reporter (*Tpm1*^*GT*^) mouse construct described in our prior publication^8^. We crossed the *Tpm1*^*GT*^ with a *FlpO* recombinase mouse (Jackson Laboratories, strain 011065) to remove the GeneTrap-Reporter cassette, leaving exon 3 flanked by loxP sites. After confirming accurate recombination by sequencing and backcrossing 3 generations on a C57BL6/J background, we crossed the *Tpm1*^*fl*^ mice with *Cdh5*^*Cre*^ mice^15^ or *Vav*^*Cre*^ mice^16^ to generate experimental mice for this study. All mice were genotyped by tail snip PCR (Transnetyx). Genotypes were additionally confirmed for experimental mice by PCR of murine whole blood using the following primers:

*Tpm1* proximal loxP Forward: AAGGCGCATAACGATACCAC

*Tpm1* distal loxP Forward: GAGGAGGCCGAGAAGGCTG

*Tpm1* distal loxP Reverse: CACAGGCTGGAGTCCCTGC

### Semiquantitative real time PCR

We confirmed excision of *Tpm1* exon 3 in the context of *Cdh5*^*Cre*^ or *Vav*^*Cre*^ by monitoring *Tpm1* mRNA expression by semiquantitative real time PCR on a QuantStudio 5 instrument (Applied Biosystems). We constructed cDNA libraries from whole mouse blood using Qiagen RNeasy kits according to the manufacturer’s instructions. We used the following primers to measure total *Tpm1* and *TATA Binding Protein (Tbp)* mRNA expression:

*Tpm1* Forward: CTGGTTGAGGAGGAGTTGGA

*Tpm1* Reverse: ATGTGCTTGGCCTCTTTCAG

*Tbp* Forward: CTCAGTTACAGGTGGCAGCA

*Tbp* Reverse: ACCAACAATCACCAACAGCA

Similar results were obtained using primers designed to specifically measure low molecular weight *Tpm1* mRNA transcripts, which we previously found to be the abundant.

### Whole mount immunohistochemistry

We set up timed matings to generate *Cdh5*^*Cre*^ *Tpm1*^*fl/fl*^ embryos and littermate controls. We harvested E9.5 embryos and genotyped tail remnants using the following primers (as also described in the genotyping strategy in Shibata *et al*^14^):

*Tpm1* proximal loxP Forward: AAGGCGCATAACGATACCAC

*Tpm1* proximal loxP Reverse: CCGCCTACTGCGACTATAGAGA

We then performed whole mount immunostaining and quantifications as previously described^8^, using antibodies to detect CD31^+^ endothelial cells (anti-mouse CD31 MEC13.3, BD Biosciences) and Runx1^+^ hemogenic cells (Runx1 monoclonal antibody EPR3099, Abcam). Secondary antibodies were Goat anti-Mouse AF488 and Goat anti-Rat AF555 (Abcam). We collected images using a Leica confocal microscope and manually quantified flat HECs using ImageJ software.

### Bone marrow isolation and analysis

Bone marrow from adult mice aged 2-3 months was isolated, stained, and analyzed as previously described^8,17^. We analyzed data using FlowJo software (v10, BD Biosciences).

### Complete blood counts

We collected blood into EDTA-coated tubes via retro-orbital collection and obtained total blood counts using a Hemavet V5 instrument (Drew Scientific).

### Genetic colocalization studies

We collected and analyzed publicly available genome wide association study (GWAS) data^1,2^ using established genetic colocalization software^19^. We inferred a PP4 > 0.8 to indicate true colocalization between two traits^5^. We generated LocusZoom plots using a web-based plotting interface (https://locuszoom.org).

### Platelet half-life experiments

We injected anti-CD42c^DyLight488^ antibody (Emfret Analytics). into *Vav*^*Cre*^ *Tpm*^*fl/fl*^ and littermate controls (*Vav*^*WT*^ *Tpm*^*fl/fl*^) and monitored the percentage of CD41^APC+^ platelets that were also CD42c^DyLight488+^ over time via serial blood collection. We performed flow cytometry on a BD Cytoflex instrument (Beckman Coulter).

### Platelet adhesion and immunofluorescence staining

We pre-treated glass cover slips with fibrinogen, fibronectin, or collagen for >2 hrs and blocked with 3% bovine serum albumin (Sigma Aldrich) for 2 h. We collected whole blood into 20 units/mL heparin and incubated on cover slips for 30 min. After washing gently with PBS, we fixed adherent cells in 4% paraformaldehyde, stained with Phalloidin-AF488 (Invitrogen) per manufacturer’s instructions, and mounted on glass slides using ProlongGold with DAPI (Invitrogen). We found that this staining strategy effectively highlighted actin-rich DAPI-negative platelets while excluding erythrocytes and rare DAPI^+^ white blood cells. We imaged slides using an Olympus XL microscope and quantified actin^+^ DAP^I-^ platelets using ImageJ.

### Platelet activation experiments

We conducted platelet activation experiments as previously described^24^. We collected whole blood via retro-orbital collection into 20 U/mL heparin (Sigma Aldrich) and activated for 10 min with indicated concentrations of Thrombin (Sigma Aldrich), Collagen Related Peptide A (CRP-A, Aniara Diagnostica). We measured αIIbβ3 integrin activation (JON/A-PE^+^, Emfret Analytics) and degranulation (CD62p/P-Selectin-FITC^+^, BD Biosciences) among CD41^+^ platelets (CD41-APC^+^, Thermo Invitrogen) by flow cytometry using a BD Cytoflex instrument (Beckman Coulter). We used FlowJo v10 (BD Biosciences) to analyze data.

### Clot contraction assay and measurement

We performed clot contraction assays as previously described^27,29,40^. Whole blood was collected directly from the inferior vena cava into syringes containing 3.2% sodium citrate as an anticoagulant at a blood-to-anticoagulant ratio of 9:1. For some experiments, platelet-rich plasma (PRP) was subsequently isolated by centrifugation of whole blood at 200g for 10 minutes. Citrated mouse blood and PRP samples were activated with 5 U/mL thrombin and 4 mM CaCl², then transferred to transparent plastic cuvettes (7×12×1 mm) prelubricated with 4% (vol/vol) Triton X-100 in 150 mM NaCl for whole blood and 1% Pluronic in 150 mM NaCl for PRP to prevent clot adhesion to the cuvette walls. Using light scatter-based tracking, changes in clot size during contraction were measured every 15 seconds over a 20-minute period. Serial images of the contracting clot were used to generate a kinetic curve, from which the extent of contraction— defined as the relative reduction in clot size after 20 minutes—was determined.

### Murine stroke model and histology

We chose to monitor the effects of ferric chloride-induced injury to the middle cerebral artery (MCA) to assess the role of platelets in this established stroke model^33^. The MCA was exposed in anesthetized mice and treated with FeCl_3_ (10% w/v, Sigma Aldrich). Laser Doppler was used to monitor MCA flow cessation following injury. The MCA clot region was then excised, sectioned, and stained for vWF and CD42d to ascertain clot morphology. Images were obtained on an epifluorescence microscope and graphical overlays were created using ImageJ.

### Tail bleeding and aPTT measurement

Tail bleeding experiments were performed as described^33^. Briefly, an incision was made in anesthetized mice and the tail submerged in 50 mL warm PBS. The time to initial cessation of bleeding, total bleeding time, amount of blood lost (as a function of murine weight), and optical density of exsanguinated blood in PBS were quantitated. We also measured activated partial thromboplastin clotting time (aPTT) in primary murine blood using standard assays^33^.

### Statistical and graphical output

Statistics were calculated using GraphPad Prism (v10) or R (v4.4). Graphical outputs used these same software packages.

## Acknowledgements

This study was supported by the National Institutes of Health (NHLBI K99 HL156052 to CST, NINDS R01 NS131279 to OAMC) and the American Heart Association (25POST1357254/2025 to ADP).

## Conflict of Interest

OAMC is a cofounder of NanoMuse, a startup company developing treatments for stroke and other neuroinflammatory conditions.

