## Supplementary Information for "Tropomyosin 1 promotes platelet adhesion and clot contraction separate from its roles in developmental hematopoiesis"

**Supplementary Figures**

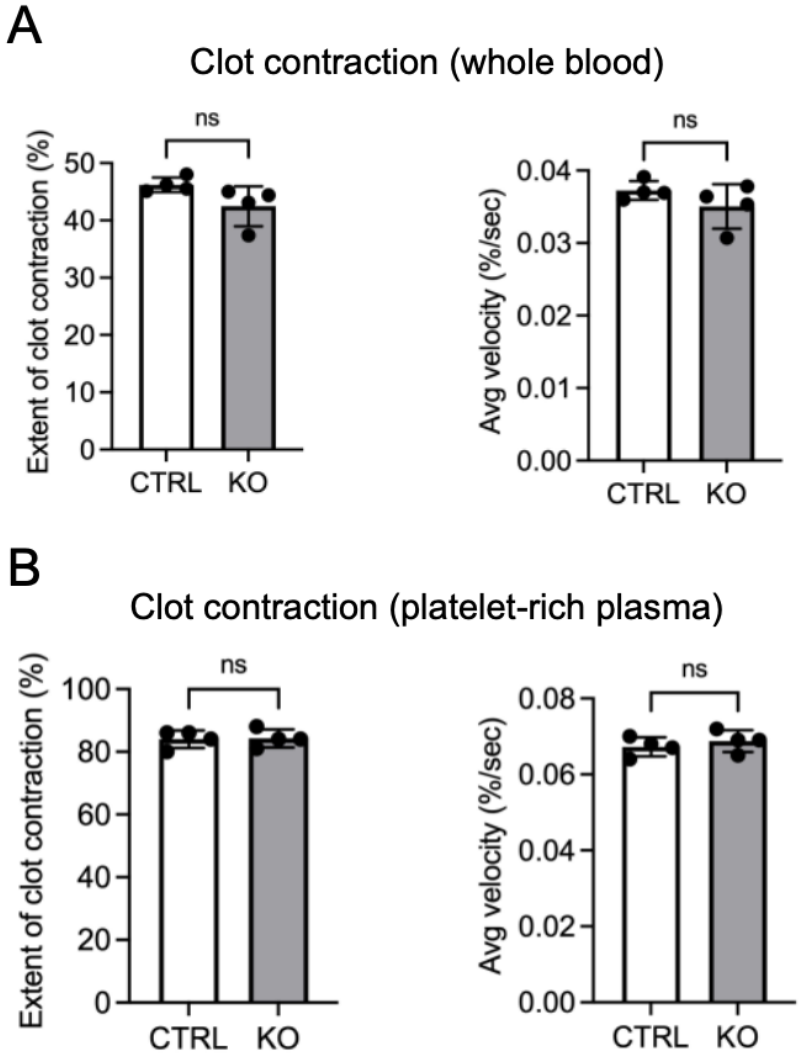


**Supplementary Figure 1. Clot contraction parameters for whole blood (WB) and platelet rich plasma (PRP).** A) Whole blood clot contraction parameters. B) PRP blood clot contraction parameters. No statistically significant changes were detected.

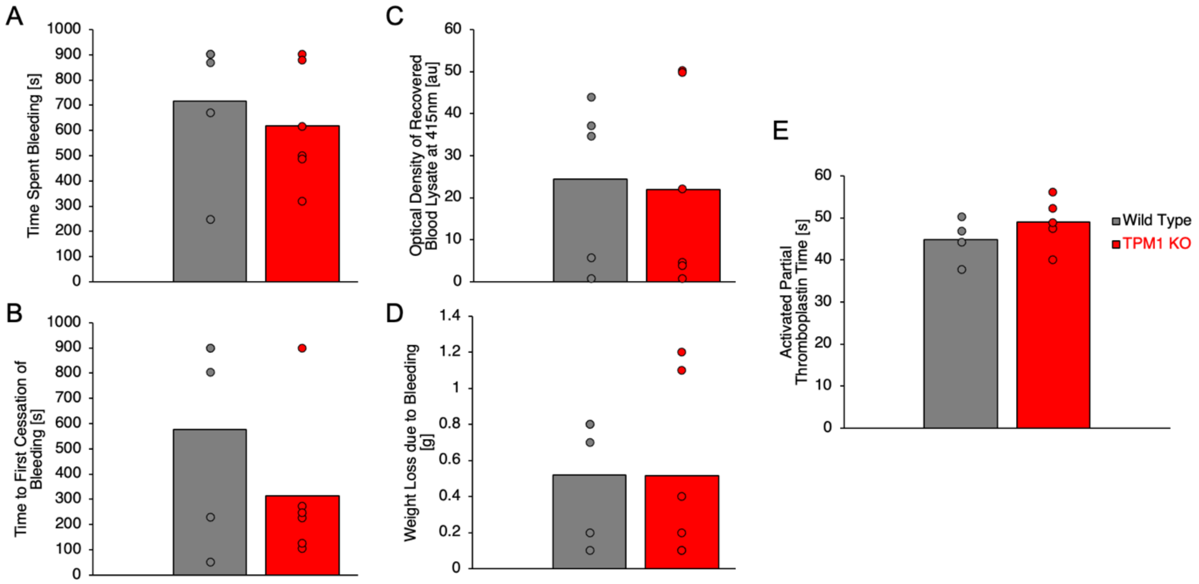


**Supplementary Figure 2. Tail bleeding and coagulation parameter measurements for *TPM1KO* and littermate controls.**
A. Time spent bleeding following tail incision.
B. Time to first cessation of bleeding.
C. Optical density of recovered blood following tail bleeding.
D. Weight of blood lost during tail bleeding.
E. Activation partial thromboplastin time. In all plots, *Vav*$^{Cre}$ *Tpm1*$^{fl/fl}$ (*TPM1KO*) are shown in red and littermate controls in gray. No statistically significant changes were detected.

### Supplementary Tables

**Supplementary Table S1.** Blood cell counts from adult *Cdh5*$^{Cre}$ *Tpm1*$^{fl/fl}$ adult mice and matched littermate controls (mean±SD for n=8-10 males per genotype). All values were within normal ranges for adult mice. Abbreviations include WBC (white blood cell count), RBC (red blood cell count), HGB (Hemoglobin), HCT (Hematocrit), MCV (mean corpuscular volume), MCH (Mean corpuscular hemoglobin), MCHC (Mean corpuscular hemoglobin concentration), PLT (Platelet count), PCT (platelet-crit), MPV (Mean platelet volume), PDW (Platelet distribution width), RDW (Red cell distribution width), LYM (Lymphocyte count), MON (Monocyte count), NEU (neutrophil count), LY% (Lymphocyte %), MO% (Monocyte %), NE% (Neutrophil %).

| Trait (units) | Control | *Cdh5*$^{Cre/+}$ *Tpm1*$^{fl/fl}$ |
|---|---|---|
| **WBC** | 8.23 (2.42) | 8.35 (5.08) |
| **RBC** | 8.05 (0.95) | 8.40 (0.42) |
| **HGB** | 12.40 (1.51) | 13.13 (0.61) |
| **HCT** | 39.06 (5.25) | 40.70 (2.15) |
| **MCV** | 56.67 (21.28) | 56.83 (21.18) |
| **MCH** | 17.98 (6.11) | 18.13 (5.64) |
| **MCHC** | 36.93 (11.69) | 37.23 (10.70) |
| **PLT** | 455.83 (54.27) | 463.83 (24.90) |
| **PCT** | 0.25 (0.03) | 0.29 (0.02) |
| **MPV** | 6.57 (2.37) | 7.18 (2.60) |
| **PDW** | 33.92 (11.91) | 34.15 (10.66) |
| **RDW** | 26.20 (9.93) | 26.77 (9.34) |
| **LYM** | 6.79 (2.81) | 6.56 (3.62) |
| **MON** | 0.24 (0.11) | 0.20 (0.15) |
| **NEU** | 1.20 (0.77) | 1.59 (1.43) |
| **LY%** | 95.63 (46.88) | 94.43 (39.49) |
| **MO%** | 3.73 (2.17) | 2.83 (1.19) |
| **NE%** | 17.32 (11.70) | 19.40 (5.64) |

**Supplementary Table S2.** Blood cell counts from adult *Vav*$^{Cre}$ *Tpm1*$^{fl/fl}$ adult mice and matched littermate controls (mean±SD for n=6 males per genotype). All values were within normal ranges for adult mice. Abbreviations include WBC (white blood cell count), RBC (red blood cell count), HGB (Hemoglobin), HCT (Hematocrit), MCV (mean corpuscular volume), MCH (Mean corpuscular hemoglobin), MCHC (Mean corpuscular hemoglobin concentration), PLT (Platelet count), PCT (platelet-crit), MPV (Mean platelet volume), PDW (Platelet distribution width), RDW (Red cell distribution width), LYM (Lymphocyte count), MON (Monocyte count), NEU (neutrophil count), LY% (Lymphocyte %), MO% (Monocyte %), NE% (Neutrophil %).

| Trait (units) | Control | *Vav*$^{Cre/+}$ *Tpm1*$^{fl/fl}$ |
|---|---|---|
| **WBC** | 7.23 (1.74) | 5.84 (1.49) |
| **RBC** | 8.13 (0.46) | 8.05 (0.78) |
| **HGB** | 13.46 (0.65) | 12.94 (1.10) |
| **HCT** | 40.58 (1.93) | 40.32 (3.52) |
| **MCV** | 50.00 (1.56) | 50.38 (2.20) |
| **MCH** | 16.60 (1.33) | 16.09 (0.54) |
| **MCHC** | 33.24 (2.68) | 32.08 (0.75) |
| **PLT** | 514.00 (72.99) | 536.57 (86.47) |
| **PCT** | 0.38 (0.17) | 0.32 (0.07) |
| **MPV** | 5.90 (0.36) | 6.14 (0.40) |
| **PDW** | 29.28 (0.92) | 30.23 (0.66) |
| **RDW** | 22.79 (4.24) | 24.13 (4.28) |
| **LYM** | 6.06 (1.41) | 4.92 (1.35) |
| **MON** | 0.15 (0.10) | 0.17 (0.06) |
| **NEU** | 1.03 (0.45) | 0.75 (0.26) |
| **LY%** | 83.92 (3.35) | 84.09 (3.48) |
| **MO%** | 2.22 (1.45) | 3.00 (1.12) |
| **NE%** | 13.85 (3.81) | 12.89 (3.18) |
